# Tau pathology reprograms glucose metabolism to support glutamatergic activity and excitatory imbalance

**DOI:** 10.1101/2025.07.25.666872

**Authors:** Riley E. Irmen, Sierra M. Turner, J. Andy Snipes, Holden C. Williams, Velmurugan G. Viswanathan, Jerry B. Hunt, Junyan Li, Patrick G. Sullivan, Daniel C. Lee, Lance A. Johnson, Shannon L. Macauley

**Affiliations:** Department of Physiology, University of Kentucky, Lexington, KY, USA; Department of Neuroscience, University of Kentucky, Lexington, KY, USA; Spinal Cord and Brain Injury Research Center, University of Kentucky, Lexington, KY, USA; Sanders Brown Center of Aging, University of Kentucky, Lexington, KY, USA

## Abstract

Alzheimer’s disease (AD) is not only characterized by amyloid-beta (Aβ) and tau pathology, but also by early and progressive disruptions in metabolism. Neuronal excitability is tightly coupled with metabolic demand, and aberrant excitatory activity – observed in AD patients and models – can drive changes in metabolism. While Aβ-related metabolic impairments are well-described, less is known about how tau pathology independently contributes to altered metabolic states and excitatory tone. Therefore, we explored how tau pathology impacted whole body and CNS metabolism in mouse models of tauopathy, including the P301S PS19 and Tau4RTg2652 mice. In both models, hyperphosphorylated tau prevents the age-related decline in whole-body metabolism by preserving glucose tolerance and mitigating shifts in fuel utilization (respiratory exchange ratio; RER), suggesting the mice are “glucose needy”. Tau pathology also preserves diurnal rhythms in hippocampal interstitial fluid (ISF) glucose and lactate, likely due to increased neuronal activity during the active (dark) phase. Stable isotope-resolved metabolomics following ^13^C-glucose administration revealed that glucose is preferentially shunted toward glutamate synthesis—at the expense of GABA—highlighting a shift in excitatory/inhibitory balance. Interestingly, these changes were not explained by a primary deficit in synaptic mitochondria but by alterations in glycolytic flux. Adaptations were time of day dependent, where ISF glutamate rises after a glucose injection in the dark period but not the light period. This suggests increased glutamatergic activity may drive metabolic demand during the dark period when mice are more active. Together, these studies fundamentally highlight the important coupling between metabolism and excitability, which is disrupted by hyperphosphorylated tau, tau aggregation, and neurodegeneration. Understanding how tau pathology and metabolism interrelate provides a novel lens for the development of therapeutic targets in late stage AD.

## Introduction

Alzheimer’s disease (AD) is the most common cause of dementia, characterized by the progressive aggregation of amyloid beta (Aβ), followed by hyperphosphorylated tau, and subsequent neurodegeneration. In addition to these pathological features, metabolic dysfunction occurs early in the AD cascade [1–11], suggesting metabolic dysfunction is not simply a downstream consequence of AD pathogenesis, but may contribute causally to disease onset and progression [12]. At the molecular level, AD pathology is associated with mitochondrial dysfunction, oxidative stress, and inflammation, which are hypothesized to underly altered energy and metabolic homeostasis [13, 14]. 2-Deoxy-2-[^18^F]fluoro-D-glucose positron emission tomography (FDG-PET), the gold standard for imaging glucose uptake in AD, has repeatedly shown cerebral hypometabolism in patients with AD and mild cognitive impairment (MCI) [15, 16]. However, it is unclear if these findings are solely due to neuronal dysfunction and cell loss, or whether Aβ or tau pathology cause shifts in brain metabolism independent of cell loss. Interestingly, recent evidence suggests that tau pathology may independently, and paradoxically, drive regional hypermetabolism in patients with 4R tauopathies or early-stage AD [17, 18]. Therefore, the relationship between tau pathology and metabolism is complex and poorly understood. These findings underscore the importance of clarifying the independent effects of tau pathology on metabolism to identify potential therapeutic targets and biomarkers of disease progression.

One mechanism possibly underlying alterations in metabolism is changes in neuronal excitability. Neuronal excitability and energy metabolism are tightly coupled and essential for maintaining healthy brain function. For instance, during periods of heightened activity, glucose uptake and oxidative phosphorylation are upregulated to sustain increased ATP production, lactate shuttling, glutamate recycling, and neurotransmitter production [19–27]. In neurodegenerative diseases, like AD, this coupling may become disrupted. Previous studies observed aberrant excitability throughout cortical and hippocampal circuits in patients and mouse models of AD, which can feed forward to exacerbate metabolic need [28–31]. However, the independent effects of tau pathology on the coupling of metabolism and excitability remain unexplored. While increased neuronal activity is known to promote tau release into the extracellular space, the extent to which tau pathology drives neuronal excitation versus silencing remains controversial [32–40]. Therefore, we sought out to investigate the interrelationship between tau pathology, metabolism, and neuronal excitability.

In this study, we characterized how tau pathology impacted the relationship between metabolism and excitability in the P301S PS19 mouse model of tauopathy. Using indirect calorimetry and glucose tolerance tests, we found that female P301S mice are resistant to age-related metabolic decline in whole body metabolism. Moreover, we found similar effects in preserved glucose tolerance using the Tau4RTg2652 mice that exhibit increased ptau levels but not neurofibrillary tangles or neurodegeneration. This data suggests that metabolic preservation is due to ptau, not tangles or neurodegeneration. Using intrahippocampal biosensors to assess interstitial fluid (ISF) glucose and lactate dynamics, a similar pattern emerged. P301S mice maintained daily ISF glucose and lactate rhythms, which are lost with age in WT mice. Using [U-¹³C]glucose stable isotope resolved metabolomics (SIRM), labeled glucose was preferentially incorporated into glutamate, which occurred at the expense of GABA and other metabolites, suggesting that glucose is an important biosynthetic substrate for glutamate production in P301S mice. Despite this, the total abundance for both glutamate and GABA were reduced in the P301S mice, highlighting tau-dependent changes in the neurotransmitter pool. To further interrogate the relationship between metabolism and excitability, P301S mice were administered a glucose challenge during the active (dark) and inactive (light) periods and changes in ISF glucose and glutamate were assessed. During the active period, P301S mice show increased glucose uptake and sustained elevations in ISF glutamate levels, suggesting metabolic driven changes in hyperexcitability. In the inactive period, ISF glutamate levels drop in WT mice, but this response is blunted in P301S mice, indicating persistent metabolic and neuronal activity during a period typically associated with inactivity and sleep. Bulk RNA sequencing of the P301S cortex demonstrated a downregulation of synaptic signaling pathways and an upregulation of metabolic and mitochondrial gene expression following a glucose challenge. Alterations in mitochondrial metabolism were not observed in synaptic mitochondria, as oxygen consumption rates (OCR) were unchanged in P301S synaptosomes compared to WT. Together, these findings suggest tau pathology causes a metabolic shift where glucose is used to simultaneously fuel glutamate production and balance hyperexcitability, resulting in a metabolically needy and inflexible system.

## Methods

### Mice

Female P301S PS19 (P301S), Tau4RTg2652 (4Rtg), and wild-type (WT; both C57B6C3F1/J mixed genetic background) littermates were used in all experiments. P301S mice develop hyperphosphorylated tau (ptau), neurofibrillary tangles, and neurodegeneration with age [41]. Mice become moribund by ∼11 months of age due to hindlimb paralysis. To avoid confounds due to hindlimb paralysis, 3, 6, and 9 month old P301S and WT mice were used in this study. 4Rtg mice overexpress wild-type human tau causing a 12-fold overexpression of tau in the brain compared to endogenous murine levels. Despite increased protein level and pre-tangle pathology by 3 months, neurofibrillary tangles are absent in the 13 month 4Rtg mice used in this study [42]. Mice were given food and water *ad libitum* and maintained on a 12:12 light/dark cycle (0600:1800). All procedures were carried out in accordance with an approved IACUC protocol from Wake Forest School of Medicine and University of Kentucky.

### Immunohistochemistry

Mice were deeply anesthetized with isoflurane and transcardially perfused with heparinized phosphate buffered saline (PBS). The mouse brains were dissected from the skull, post-fixed in 4% paraformaldehyde (PFA) for 48 hours at 4°C, and then cryoprotected in 30% sucrose at 4°C. Sectioning was performed on a freezing microtome at 40µm and were stored in cryoprotectant until use. Serial sections through the anterior-posterior aspect of the hippocampus were immunostained for hyperphosphorylated tau using a biotinylated AT8 monoclonal antibody (anti-tau pSer202, Thr205, Thermofisher, #MN1020B, 1:) [43, 44]. AT8 immunostaining was developed using a Vectastain ABC kit (PK-6100, Vector Labs) and DAB reaction (ICN 980681, Fisher Scientific) and imaged using a Zeiss Axio Scan.Z1 slide scanner (Carl Zeiss Microscopy GmbH, Jena, Germany). For quantification, scanned images were converted to 16-bit grayscale and thresholded to highlight AT8 staining. Percent occupied by AT8 (e.g. mean reactivity) and region volume/thickness were quantified by a blinded researcher throughout the entorhinal cortex, hippocampus, and cortex.

### Glucose tolerance tests

3, 6, and 9 month old P301S, 4Rtg, and WT mice were fasted for four hours and then a baseline fasted blood glucose level was measured using a glucometer (Contour, Bayer). Mice were given a glucose bolus (2g/kg) via intraparietal (i.p.) injection as previously described [45]. Blood glucose levels were measured at 15, 30, 45, 60, and 90 minutes post injection via tail bleed.

### Peripheral metabolic characterization

Indirect calorimetry, feeding behavior, and activity were measured using the TSE Phenomaster system (TSE System Inc.). Oxygen consumption (VO2), carbon dioxide production (VCO2), total energy expenditure (TEE), respiratory exchange ratio (RER), locomotor activity, and food intake were measured for 72 hours following a 24-hour acclimation period. Mice were housed individually in recording chambers and maintained on a 12:12 light/dark cycle with food and water *ad libitum* for the duration of the experiment.

### Guide cannula surgery

Mice were anesthetized with 5% isoflurane prior to surgery and maintained at 1.5-2.5% isoflurane during surgery. Sterile field was maintained to prevent infection during surgery. Guide cannulas (MD-2255 BASi Research Products) were stereotaxically placed and implanted bilaterally in the hippocampi (from bregma, A/P −3 mm, M/L +/− 3 mm, D/V −1.8 mm) as previously described [46]. Mice were placed into sampling cages (Pinnacle Technology) and allowed to recover for 72 hours after surgery prior to EEG recording and maintained on a 12:12 light/dark cycle.

### Biosensor recording and analysis

Prior to implantation, biosensors were tested for response specificity to each analyte (L-Lactate, D-Glucose, L-Glutamate) and ensured no response to interference analyte (ascorbic acid). 24 hours post-surgery, mice were briefly anesthetized with isoflurane for the implantation of two amperometric biosensors, specific to either glucose, lactate, or glutamate (7004, Pinnacle Technology) into the guide cannula placed during surgery. Biosensors were connected to a flexible preamplifier (100x amplification) and attached to the mouse’s headmount. The preamplifier was connected to a commutator, which passes electrical signal to the data acquisition system (8401, Pinnacle Technology) for sampling at 1 Hz for biosensor channels. This set-up allows for unanesthetized, unrestrained movement throughout the experiments. 72-hour diurnal rhythm recordings began after a stable baseline reading was reached using Sirenia Acquisition software. All biosensor data was exported from Sirenia software, combined into 10 second bins, and converted from nA to mM using the calibration constant for that biosensor. Binned biosensor data was then analyzed as fluctuations over the diurnal day and light/dark periods.

### Mitochondrial bioenergetics

9 month WT and P301S brains were rapidly extracted. Mitochondrial bioenergetics were measured using the Seahorse XFe96 Extracellular Flux Analyzer (Agilent Technologies, USA) as previously established [46–49]. Synaptic and non-synaptic mitochondria were diluted in mitochondrial buffer (MRB) (125 mM KCl, 0.1% BSA, 20 mM HEPES, 2 mM MgCl2, and 2.5 mM KH2PO4, adjusted pH 7.2 with KOH). A total of 3 µg of synaptic and 1.25 µg of non-synaptic mitochondria were loaded per well. Oxygen consumption rates (OCR) were assessed under various respiratory states using substrates, inhibitors, and uncouplers of the electron transport chain, as per a lab-optimized protocol. Data were collected through sequential injections, mixing, equilibrium, and OCR measurements. Outputs included State III respiration (pyruvate, malate, ADP), State IV respiration (oligomycin), uncoupled respiration complex-I (FCCP), and uncoupled respiration complex-II (rotenone/succinate).

### ^13^C-glucose oral gavage and tissue collection for stable isotope resolved metabolomics

9 month old P301S and WT mice were single housed and fasted for four hours. Mice were given an oral gavage of 250 µL of 32 mg/ml ^13^C-glucose (Cambridge Isotope Laboratories Inc. #110187-42-3) as previously described [46, 50]. Briefly, 45 minutes after oral gavage, mice were cervically dislocated, brain was rapidly extracted, and flash frozen in liquid nitrogen. Stable Isotope resolved metabolomics (SIRM) and gas chromatography mass spectrometry (GCMS) were performed on one hemisphere from brain as previously described [50].

### Bulk RNA sequencing

RNA isolation was performed on the posterior cortex from the ^13^C-glucose oral gavage 9 month P301S and WT mice as previously described [51]. Briefly, 500 µL of trizol was added to cortex in centrifuge tube, homogenized using plastic pestle for 1 minute, and were allowed to sit for 5 minutes. 100 µL of chloroform was added to each sample and vigorously shaken for 15 seconds then allowed to sit for 3 minutes. The samples were spun at 12000xg for 10 minutes at 4° C. Chloroform was removed, 200 µL of 70% ethanol was added to samples, and samples purified using RNeasy columns and reagents (QIAGEN) per manufacturer’s instructions. RNA concentration was measured by loading 2 µL of samples onto a Take3 Microplate and RNA measurements were made using a Biotek Synergy H1 microplate reader using the Gen5 Take3 software module. Isolated RNA samples were then sent for bulk RNA sequencing (Novogene Co., Ltd) for sample quantitation, integrity, and purity using Agilent 5400. DESeq2 was used to determine differentially expressed genes (DEGs) using Partek Flow Software. Enrichr was used for pathway analysis of DEGs. SR Plot was used for data visualization.

### Jonckheere-Terpstra-Kendall (JTK) analysis

Biosensor recording data was preprocessed to average groups values for each Zeitgeber Timepoint. JTK analysis was done using R package MetaCycle [46]. Period measured the average cycle length in time of each rhythm. Lag indicates ZT of peak. Amplitude indicates the average difference between peak and trough each rhythm.

### Statistical analysis

All of the statistical analysis was done using GraphPad Prism 10 (GraphPad Software, LLC) or Partek Flow Software. A p-value of p<0.05 was used for determining statistical significance. For all groups, Grubb’s test was used to determine and remove outliers (alpha = 0.05). All data is reported as means with +/− SEM. Specific statistical tests used were as follows:

Figure 1: Two-way ANOVA with Tukey’s correction for multiple comparisons.
Figure 2: Area Under the Curve; Two-way ANOVA with Tukey’s correction for multiple comparisons.
Figure 3: Two-way ANOVA with Tukey’s correction for multiple comparisons.
Figure 4: Two-way ANOVA with Tukey’s correction for multiple comparisons.
Figure 5: Two-way ANOVA with Tukey’s correction for multiple comparisons.
Figure 6: unpaired t-tests; DESeq2 was used to determine differentially expressed genes using Partek Flow Software (p<0.05). Upregulated genes displayed fold change >1 and downregulated genes <-1. Enrichr was used for pathway analysis. SR Plot was used for data visualization.
Figure 7: Two-way ANOVA with Sidak’s correction for multiple comparisons.
Figure 8: Area Under the Curve (net change from baseline); unpaired t-tests.
Supplemental Figure 1: Two-way ANOVA with Tukey’s correction for multiple comparisons.
Supplemental Figure 2. Unpaired t-tests, Two-way ANOVA with Sidak’s correction for multiple comparisons, Area Under the Curve.
Supplemental Figure 3: Two-way ANOVA with Tukey’s correction for multiple comparisons.
Supplemental Table 1: Jonckheere-Terpstra-Kendall analysis.
Supplemental Figure 4: Two-way ANOVA with Sidak’s correction for multiple comparisons.
Supplemental Figure 5: Area Under the Curve (net change from baseline); unpaired t-tests.

## Results

### Tau pathology decreases cortical and hippocampal volume

As P301S mice age, AT8+ tau pathology is observed in the somatosensory/parietal/retrosplenial cortex, hippocampus, and entorhinal/piriform cortex by 6 months of age and progresses with age (Fig 1A, B). By 6 months, AT8+ staining is increased by 15% in the entorhinal/piriform cortex (Fig 1B; p<0.01) as well as by 5% somatosensory/parietal/retrosplenial cortex and hippocampus (Fig 1B; p<0.05). Volumetric analysis of the cortex and hippocampus demonstrated a 4% reduction in somatosensory/parietal/retrosplenial cortical volume as early as 6 months (Fig 1C; p<0.05) while entorhinal/piriform and hippocampal volume were decreased by 9 months (Fig 1C, p<0.05). Together, hyperphosphorylated tau (ptau) pathology exhibits region specific accumulation, which is associated with cortical atrophy as early as 6 months of age. Both tau pathology and regional atrophy progress with age.

**Figure 1.**
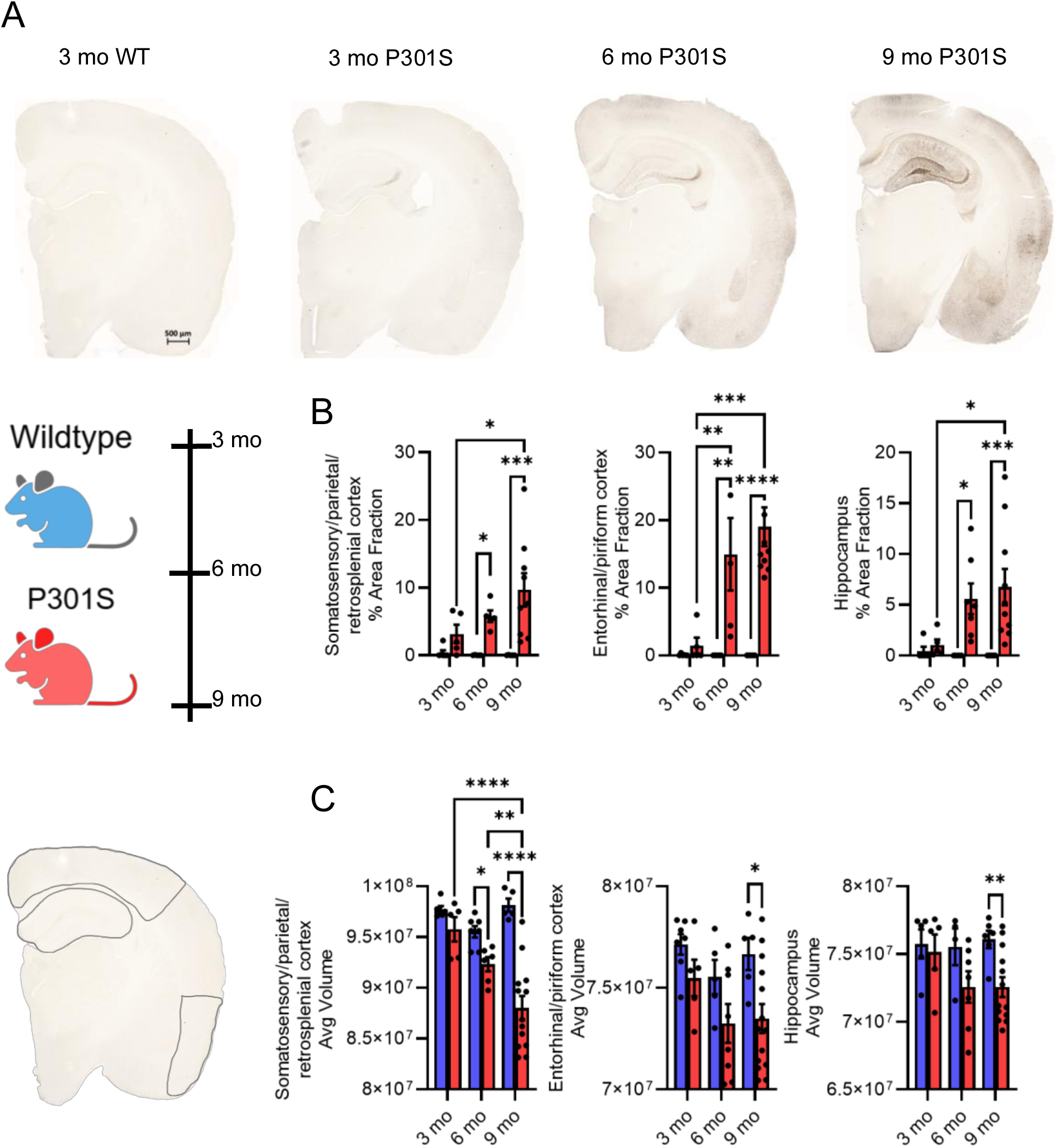
Tau pathology decreases cortical and hippocampus volume. (A) Representative images of AT8+ tau pathology in 3, 6, and 9 month P301S v. 3 month WT mice. (B) P301S (red) mice exhibit increased AT8 staining within the somatosensory, parietal, and retrosplenial cortices and hippocampus at 6 and 9 months compared to age matched WT (blue), or 3 month P301S mice. AT8+ staining is higher in entorhinal/piriform cortex compared to other cortices (somatosensory, parietal, or retrosplenial cortices). (C) Cortical atrophy begins at 6 months in P301S mice compared to aged matched WT. This atrophy continues to progress at 9 months in P301S mice. Hippocampal and entorhinal/piriform cortex atrophy is present in 9 month P301S mice. *n =* 5-10 mice/group. Statistical significance was determined using a two-way ANOVA with Tukey’s post-hoc tests. Data is represented by means ± SEM. *p<0.05, **p<0.01, ***p<0.001, ****p<0.0001

### Tau pathology preserves peripheral glucose sensitivity

To analyze whole body metabolism relative to normal aging and tau pathology, we utilized glucose tolerance tests (GTT) to examine glucose sensitivity and tolerance. Fasting blood glucose levels did not differ at baseline based on either age or genotype (Fig 2A). Wildtype (WT) mice display an age-related change in glucose sensitivity, including a higher glucose peak and delayed return to baseline following an i.p. glucose challenge (Fig 2B, p<0.001). WT mice also exhibit an age-dependent increase in area under the curve (AUC) (Fig 2C, p<0.01) during the GTT, suggesting glucose intolerance. In contrast, P301S mice age do not display an age-related change in glucose tolerance or sensitivity (Fig 2C). During the GTT, the glucose peak, return to baseline, and AUC do not change in P301S mice as they age and remains comparable to the phenotype of 3 month WT (Fig 2). This suggests glucose tolerance is preserved in 9 month P301S mice compared to 9 month WT mice (Fig 2C, p<0.01). Next, we explored if changes in body weight were driving changes in peripheral metabolism. WT mice exhibit a progressive increase in body weight with age (Suppl. Fig 1, p<0.0001); however, P301S mice weigh less at 9 months than to 9 month WT littermates (Suppl. Fig 1A, p<0.0001). Interestingly, this difference in weight gain is not due to changes in food intake (Suppl. Fig 1B). To explore whether this was due to the presence of ptau, neurofibrillary tangle formation, or neurodegeneration, we performed GTTs on 13 month Tau4RTg2652 (4Rtg) mice that exhibit elevations in ptau but no neurofibrillary tangles or neurodegeneration [42]. We found the 4Rtg mice exhibit decreased body weight and preserved glucose tolerance (Suppl. Fig 2), similar to the P301S mice. Together, this data suggests that elevations in hyperphosphorylated tau-not aggregation or neurodegeneration-drives changes in peripheral metabolism and prevents age-dependent glucose intolerance and weight gain.

**Figure 2.**
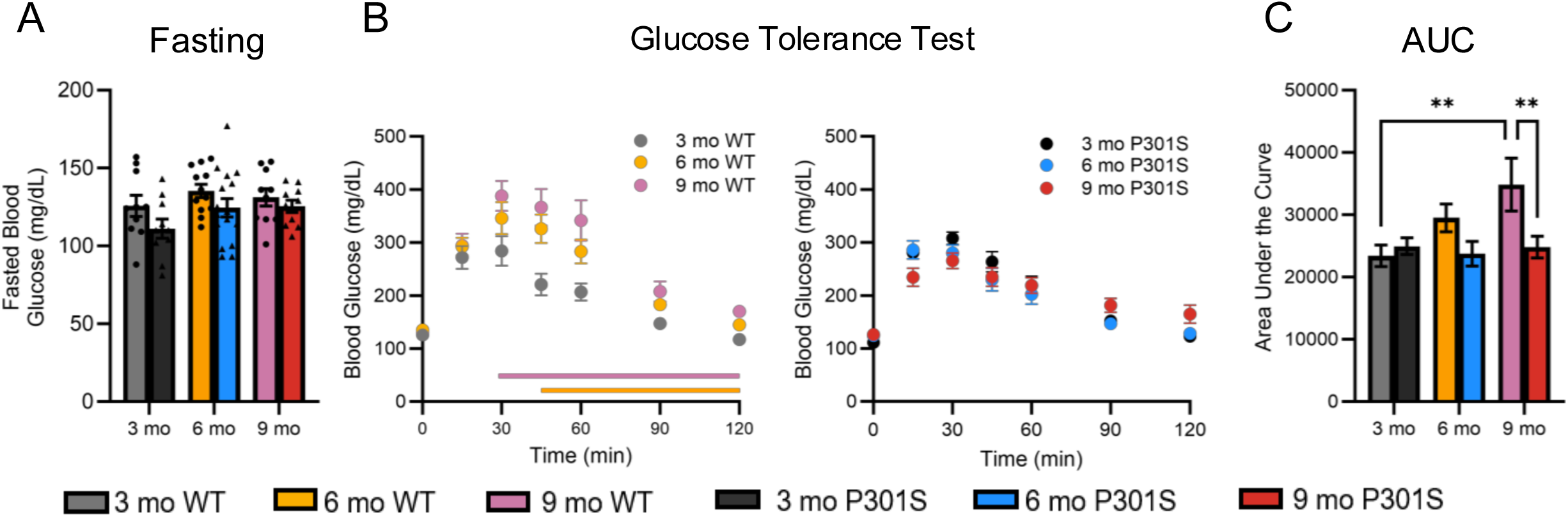
Tau pathology prevents age-related peripheral glucose intolerance. (A) P301S and WT exhibit no changes in fasting blood glucose with age. (B) As WT mice age from 3-9 months, they become glucose intolerant as evidenced by an increased peak and longer tail on glucose tolerance tests (GTTs). Orange and pink lines indicate elevated blood glucose in 6 and 9 month WT mice (respectively) compared to 3 months. Conversely, tau pathology preserves peripheral glucose tolerance independent of age. (C) Area under the curve (AUC) increases between 3 - and 9-month WT mice, while P301S have a decrease in AUC at 9 months compared to WT mice. Data reported as means ± SEM. *n =* 9-12 mice/group. Significance determined using two-way ANOVA with Tukey’s post-hoc corrections. *p<0.05, **p<0.01, ***p<0.001, ****p<0.0001

### Tau pathology slows age-dependent decreases in respiratory exchange ratio and locomotor activity

Given the tau-dependent preservation of glucose tolerance, we used indirect calorimetry to further assess how aging and tau pathology differentially affected whole body metabolism (Fig 3). We found that total energy expenditure did not change between age-matched P301S and WT mice, despite increased oxygen consumption (VO2) and carbon dioxide production (VCO2) in 9 month P301S mice (Suppl. Fig. 3A and 3B). Since this suggests metabolic insufficiency or changes in fuel preference, we calculated the respiratory exchange ratio (RER) as a measure of whole body fuel utilization. A respiratory exchange ratio (RER) value closer to 1 indicates carbohydrate (glucose) utilization and a value closer to 0.7 indicates fatty acid oxidation (FAO). As WT and P301S mice age, RER is decreased in 9 month old mice compared to 3 or 6 month genotype-matched mice (Fig 3A, WT: p<0.001, P301S: p<0.05) across the 24 hour cycle, suggesting a switch in fuel preference from carbohydrates to fats with age. However, 9 month P301S mice exhibit an increased RER compared to 9 month WT (Fig 3A, p<0.05), demonstrating that P301S mice display a preference for carbohydrates over fats when compared to age matched WT littermates. Interestingly, this was driven by RER changes in the dark period when mice are active. Therefore, we explored locomotor activity across the circadian day in P301S and WT mice relative to age. Here, we found that 9 month P301S mice are more active than their age-matched WT littermates, but only during the dark (active) period (Fig 3B, p<0.05). Together, these data suggest tau pathology preserves whole body metabolism, with a preference towards increased carbohydrate utilization and locomotor activity during the dark period in response to tauopathy.

**Figure 3.**
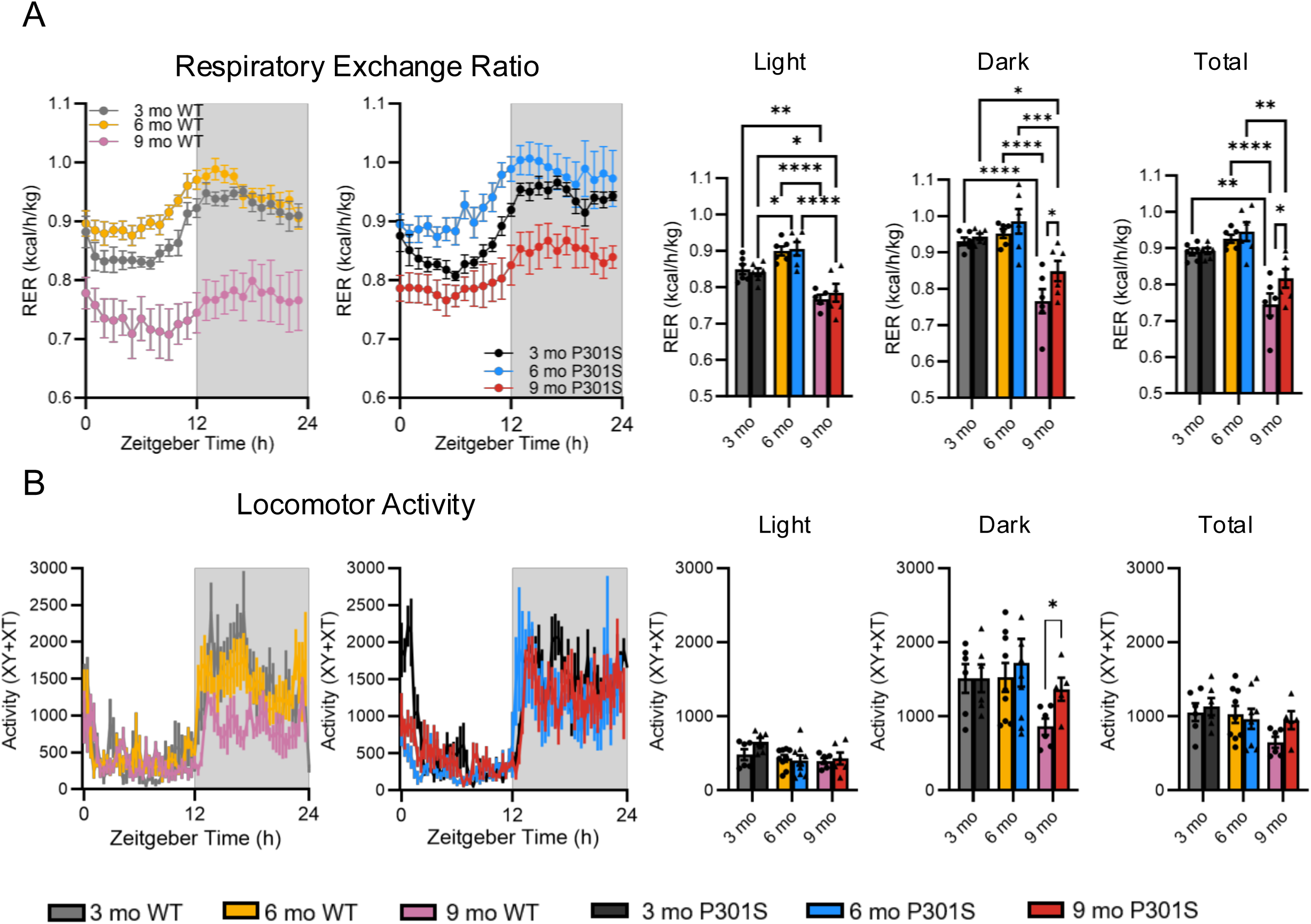
Tau pathology slows age-dependent decreases in respiratory exchange ratio and locomotor activity during the dark or active period. (A) There is an aged-related decrease in respiratory exchange ratio (RER) across the 24hr day in WT mice. P301S mice, conversely, exhibit an increase in RER from 3 to 6 months during the light period. At 9 months, RER is elevated during the dark period and across the 24hr day in P301S mice compared to WT, indicating increased carbohydrate utilization. (B) Locomotor activity in 9 month P301S mice is increased during the dark period only compared to age matched WT. Data reported as means ± SEM. *n =* 6-9 mice/group. Significance determined using two-way ANOVA with Tukey’s post-hoc correction. *p<0.05, **p<0.01, ***p<0.001, ****p<0.0001

### Tau pathology preserves hippocampal ISF glucose and lactate rhythms

Given that tau pathology shifts whole body metabolism in a manner different from normal aging, we next investigated how tau pathology and aging impact brain metabolism relative to time of day. Using intrahippocampal biosensors, we measured interstitial fluid (ISF) glucose and lactate levels across the circadian day. Young, healthy WT mice display a diurnal rhythm in ISF glucose and lactate, where both ISF glucose and lactate levels peak at ZT16-17 during the active or dark period and drop during the light period when mice traditionally sleep (Fig 4&5). By 9 months, WT mice lose fluctuations in ISF glucose levels across the light and dark period, with an accompanying reduction in rhythm amplitude (Fig 4A, C, p<0.05). While Jonckheere-Terpstra-Kendall (JTK) analysis confirmed rhythmicity in WT mice, 9 month WT mice exhibit 90% amplitude reduction and peak during the light period suggesting loss of rhythmicity and phase shift. (Fig 4C, D, Suppl. Table 1A). P301S mice, on the other hand, maintain robust ISF glucose rhythms across the light and dark periods (Fig 4B, C, p<0.0001), where both rhythm amplitude and phase are maintained as confirmed by JTK analysis (Suppl. Table 1). This data suggests aging dampens ISF glucose rhythms, while tau pathology preserves rhythmicity.

**Figure 4.**
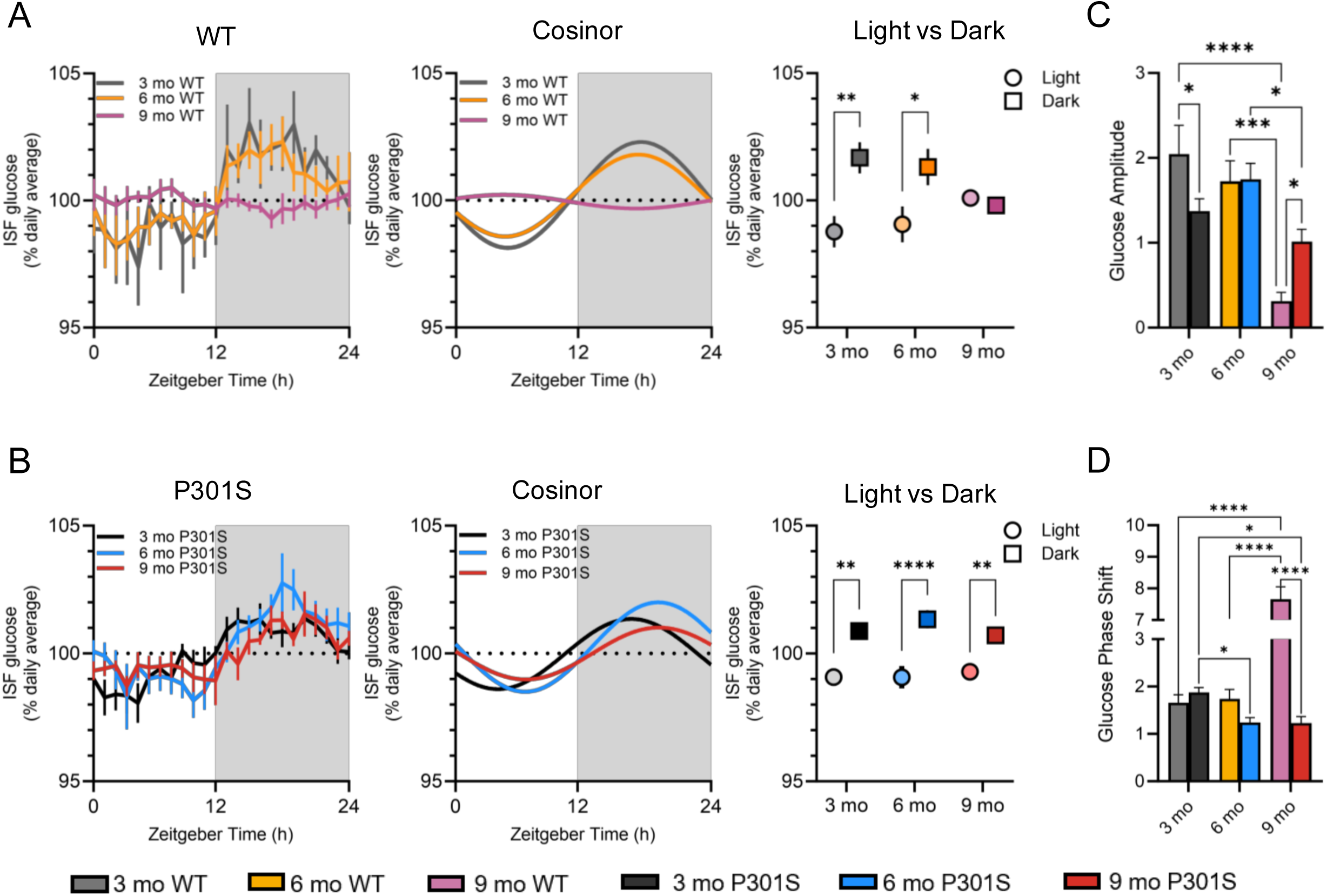
Tau pathology preserves interstitial fluid (ISF) glucose rhythms. (A) At 3 and 6 months old, WT mice exhibit robust glucose rhythms that peak during the dark period and trough in the light period. These light/dark differences are lost by 9 month in WT mice. (B) P301S mice exhibit robust diurnal ISF glucose rhythms with differences between the light and dark periods at 3, 6, and 9 months. (C) 9 month WT mice exhibit decreased ISF glucose rhythm amplitude compared to 3 and 6 month WT mice. Conversely, the amplitude of 9 month P301S mice is maintained despite a lower amplitude rhythm at 3 months of age. (D) 9 month WT mice exhibit a phase shift, or difference in rhythm peak, compared to other groups, highlighting the age-dependent change in ISF glucose rhythms. This effect was not observed in 9 month P301S mice. Data reported as means ± SEM. *n =* 6-10 mice/group. Significance determined using two-way ANOVA with Tukey’s post-hoc correction. *p<0.05, **p<0.01, ***p<0.001, ****p<0.0001

**Figure 5.**
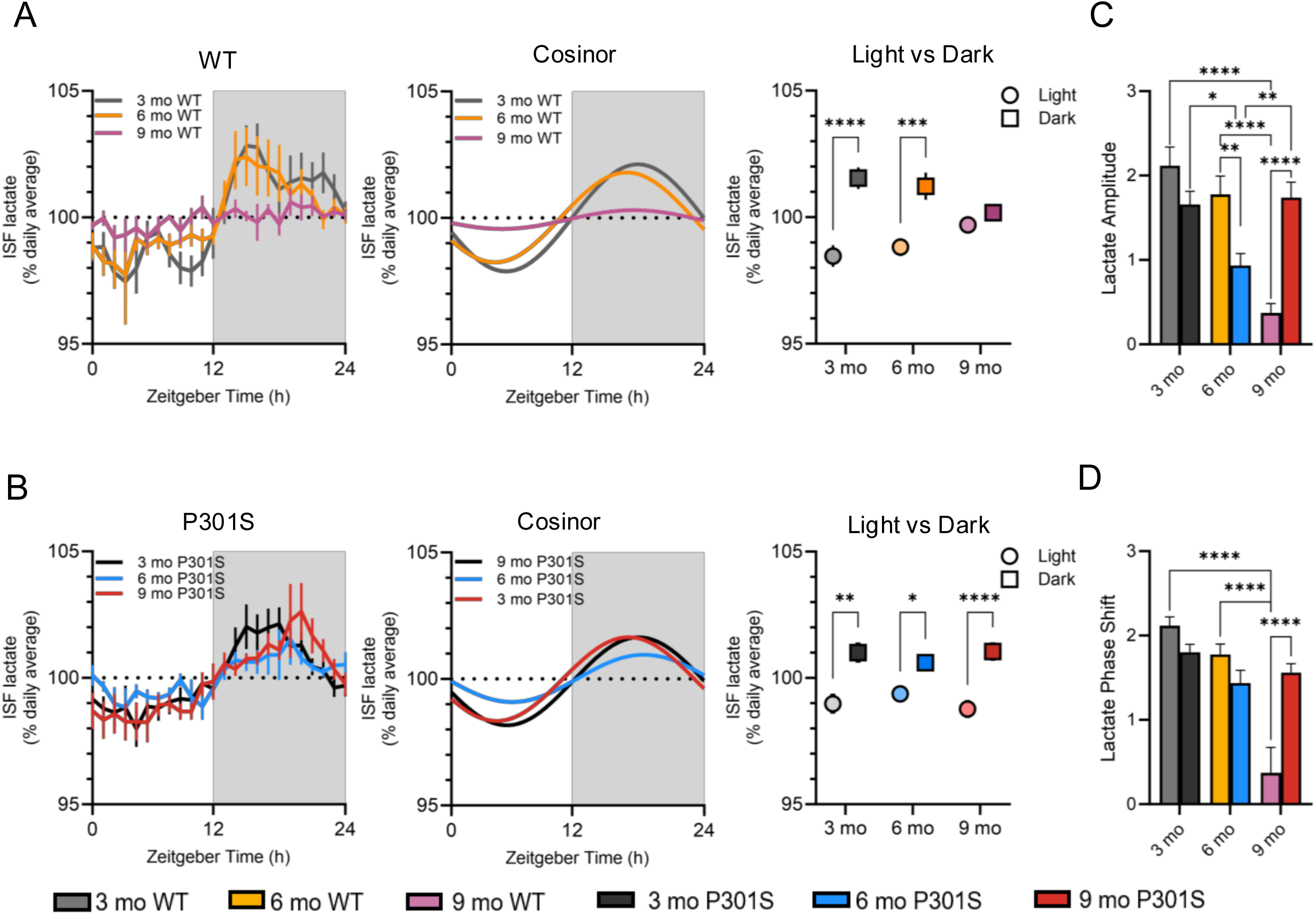
Tau pathology potentiates interstitial fluid (ISF) lactate rhythms. (A) At 3 and 6 months, WT mice exhibit robust, diurnal lactate rhythms shown by peaks in the dark period and trough in the light period. These light/dark differences are lost by 9 month in WT mice. (B) P301S mice exhibit robust diurnal ISF lactate rhythms with differences between the light and dark periods at 3, 6, and 9 months. (C) 9 month WT mice exhibit decreased ISF lactate rhythm amplitude compared to younger WT mice, while P301S mice have a potentiated rhythm at 9 months. (D) Phase shift, or peak amplitude, in 9 month WT mice differs when compared to younger WT mice. This was not observed in 9 month P301S mice. Data reported as means ± SEM. *n =* 6-10 mice/group. Significance determined using two-way ANOVA with Tukey’s post-hoc correction. *p<0.05, **p<0.01, ***p<0.001, ****p<0.0001

Similar effects were observed with diurnal fluctuations in ISF lactate. 9 month WT mice exhibit decreased ISF lactate rhythm shown by a loss in light-dark fluctuations, decreased rhythm amplitude, and loss of rhythms using JTK analysis (Fig 5A, C, Suppl. Table 1B). In contrast, tau pathology results in the maintenance of ISF lactate rhythmicity at 9 months in P301S mice, where light-dark fluctuations and rhythm amplitude are preserved (Fig 5A, C, Suppl. Table 1). Together, these findings show that tau pathology prevents age-related loss of ISF lactate rhythms. In fact, ISF lactate rhythms are potentiated in P301S mice compared to WT mice at 9 months. These findings suggest tau pathology may reprogram systemic and brain metabolism in a way that resists age-related metabolic decline or promotes metabolic inflexibility.

### Increased glucose utilization is necessary for glutamate production in response to tau pathology

Given that tau pathology preserves brain and peripheral metabolic function, we next used ^13^C-glucose stable isotope resolved metabolomics (SIRM) coupled with gas chromatography mass spectrometry (GCMS) to determine how glucose is metabolized by the brain and what biosynthetic pathways glucose supports (Fig 6A). A heatmap of relative abundance for each ^13^C-glucose labeled metabolite measured in individual mice and averaged across genotypes for 9 month P310S and WT brains is shown in Figure 6B. While the heatmap suggests that overall, the P301S brain exhibits increased ^13^C enrichment following a ^13^C-glucose challenge than a WT brain, we only observed enrichment for ^13^C-labled glutamate in P301S mice compared to controls (Fig 6B, p<0.01). This occurred at the expense of ^13^C-labeled GABA, which was decreased in P301S mice compared to WT mice (Fig 6B, p<0.05). Interestingly, heatmaps of individual and group averaged total abundance (e.g. labeled + unlabeled pools) suggests the opposite-that total pool size of these central carbon metabolites was generally lower in the P301S compared to WT, including a significant decrease in GABA (Fig 6C, p<0.05) and trending decrease in glutamate (Fig 6C, p=0.0527). These data suggest that the brain redirects glucose for glutamate biosynthesis in presence of tau pathology, rather than for traditional energy producing pathways, like glycolysis or oxidative phosphorylation. It further suggests that changes in whole body glucose metabolism could be driven by excitatory/inhibitory imbalance in the tauopathy brain.

**Figure 6.**
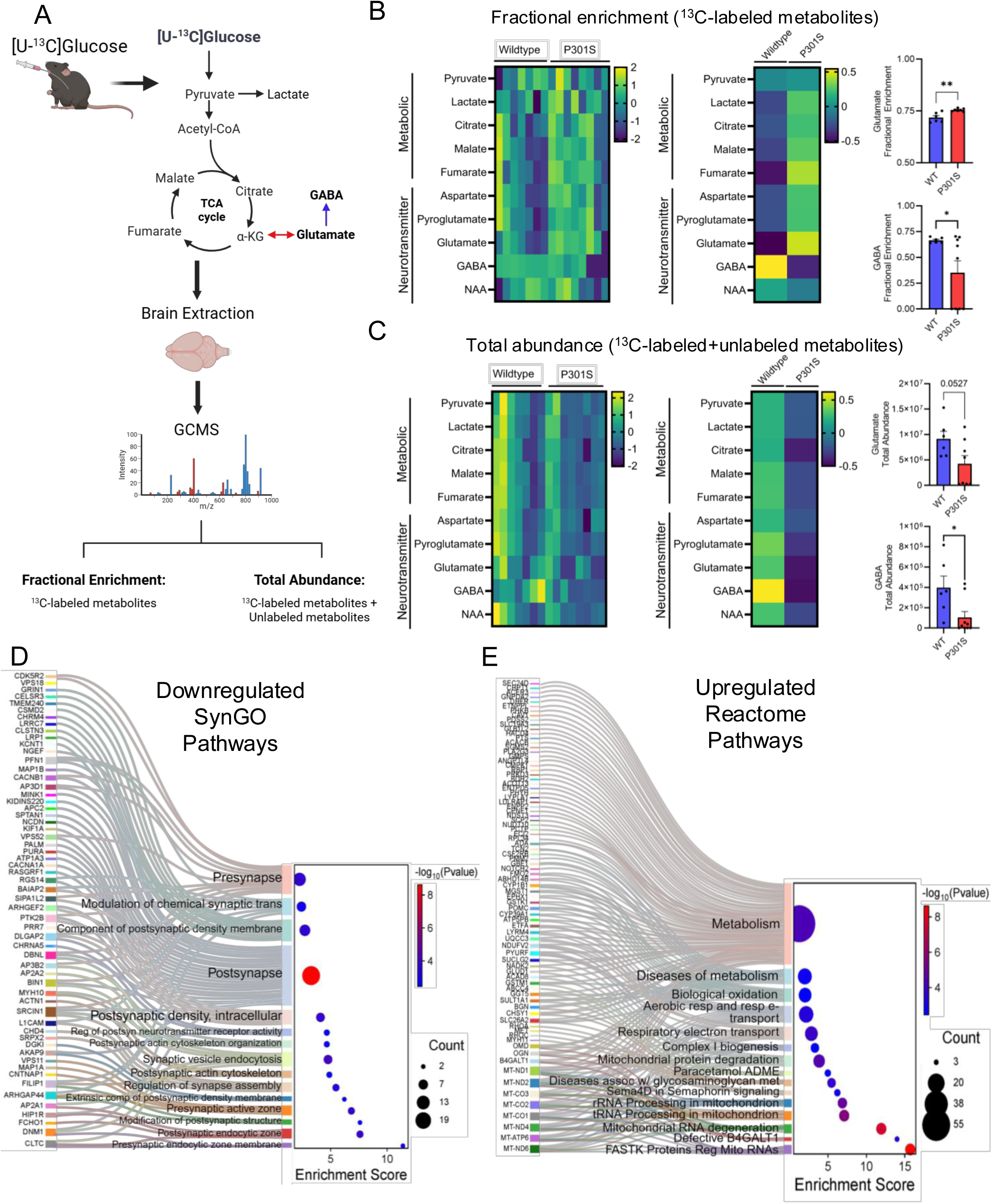
Tau pathology shunts glucose towards glutamate production while upregulating metabolic pathways. (A) Experiment design for stable isotope resolved metabolomics (SIRM) following a ^13^C-labeled glucose oral gavage. (B) Heatmap of metabolites that are fractionally enriched following ^13^C-glucose administration in individual mice and genotype averages in 9 month P301S v WT mice. When quantified, there was increased ^13^C-labeled glutamate P301S brains compared to WT. Concurrently, there was a decrease in ^13^C-labeled GABA in P301S mice compared to WT. (C) Heatmap of the total abundance (^13^C-labeled and unlabeled pools) of metabolites following ^13^C-glucose gavage. The total abundance of GABA is decreased in 9 month P301S mice compared to WT. (D,E) Gene pathway analysis following bulk RNA sequencing of 9 month P301S and WT cortical tissue post ^13^C glucose gavage yielded changes in excitatory-metabolism coupling. (D) SynGO pathways, including those involved in pre- and postsynaptic homeostasis, are downregulated in the P301S cortex compared to WT, as visualized by Sankey and dot plots. (E) Reactome pathways associated with metabolism, diseases of metabolism, and mitochondrial homeostasis are upregulated in in the P301S cortex compared to WT, as visualized by Sankey and dot plots. Data reported as means ± SEM. *n = 7*-8 mice/group. Significance determined using unpaired t-test. Enrichr was used for pathways analysis of DEGs. SR Plot was used for data visualization. *p<0.05, **p<0.01

Gene pathway analysis of the cortical transcriptome following ^13^C-glucose administration revealed interesting differences between 9 month P301S and WT mice related to synaptic and metabolic activity. First, P301S mice exhibit decreased expression of genes associated with Synapse Gene Ontologies, or SynGO, pathways. Specifically, there was decreased enrichment of pathways associated with synaptic homeostasis, synaptic transmission, synaptic density, and synaptic structure (Fig 6D). While this is most likely due to the neurodegeneration that accompanies tau pathology (Fig 1), it is striking given the increased labeled glutamate that occurs following ^13^C-glucose administration. We also noted increased enrichment of genes associated with Reactome pathways after glucose administration. Specifically, increased enrichment of genes associated with metabolism, diseases of metabolism, aerobic respiration, respiratory electron transfer, complex I biogenesis, and other pathways associated with mitochondrial processes (Fig 6E). Taken together, these data suggest that tau pathology utilizes glucose to fuel metabolic neediness in response to neuronal dyshomeostasis.

### Oxygen consumption rate is maintained in synaptic mitochondria in P301S mice

Given that complex impact of tau pathology on metabolic and neuronal function, we next examined whether primary mitochondrial dysfunction occurs in synaptosomes from P301S mice. To do this, we quantified mitochondrial respiration in synaptic and non-synaptic (neuronal soma, glia, vascular cells) mitochondria isolated from the 9 month P301S and WT brains. We noted no differences in oxygen consumption rate (OCR) in synaptic mitochondria (Fig 7A), suggesting synaptic mitochondrial function was intact in P301S mice. In non-synaptic mitochondria, we found a tau-dependent decrease in State V (succinate) compared to age-matched controls (Suppl. Fig 4A, p<0.05), suggesting impaired complex II activity. These data suggest that tau pathology does not drive primary deficits in oxygen consumption in synaptic mitochondria, but may cause metabolic inefficiency in the neuronal soma, glia, or both. It further suggests that primary mitochondrial deficits do not appear to drive the metabolic adaptations observed in P301S mice.

**Figure 7.**
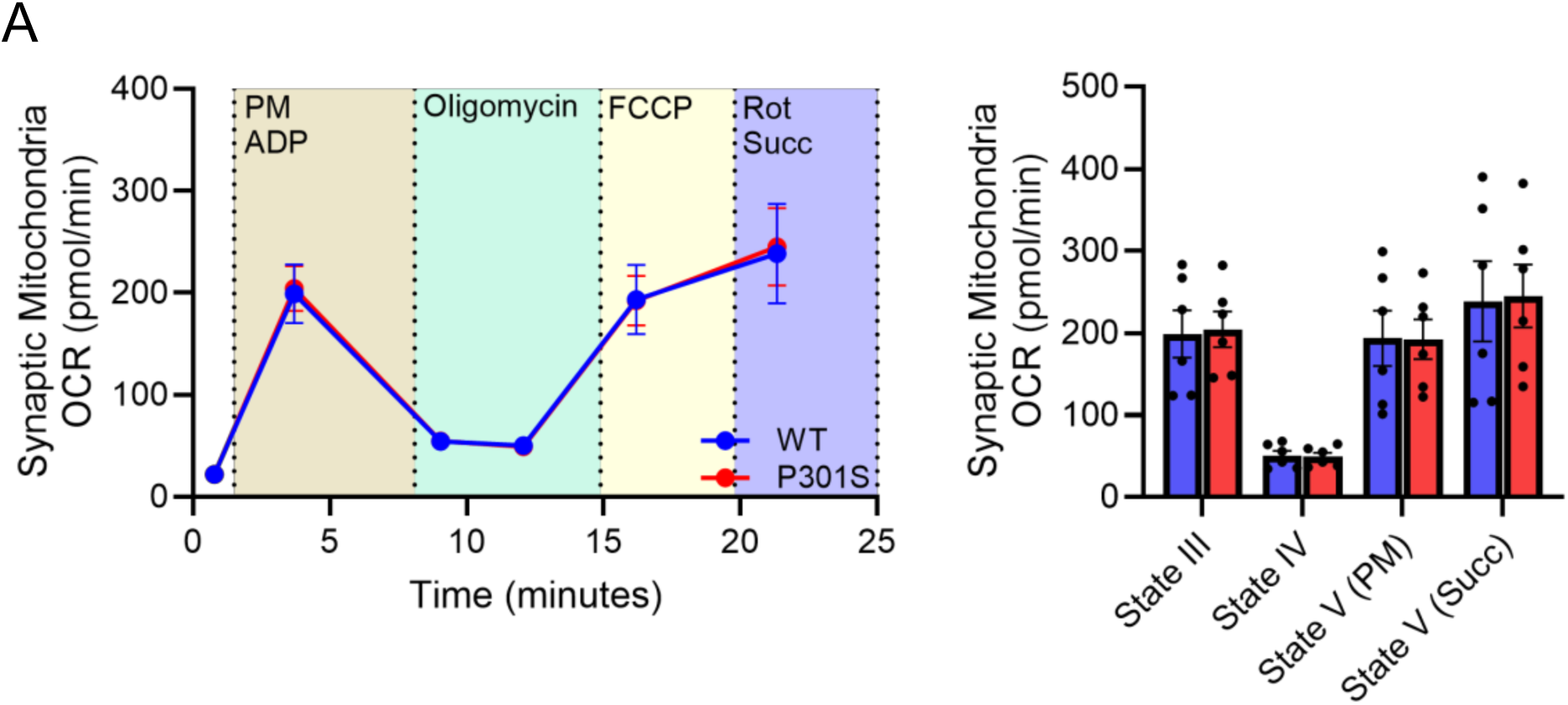
Oxygen consumption rate is maintained in synaptic mitochondria with tau pathology. (A) P301S mice show no alterations in mitochondrial oxygen consumption (OCR) in isolated synaptosomes compared to WT controls. Data reported as means ± SEM. *n =* 6 mice/group. Significance determined using two-way ANOVA with Sidak’s post-hoc correction.

### Glucose fuels ISF glutamate levels in P301S mice

Given that tau-dependent changes in glucose utilization seem to be affected by time of day, we next examined how ISF metabolites respond to a peripheral glucose challenge during the light versus dark phase. ISF glucose or glutamate biosensors were implanted into the hippocampus of 9 month P301S mice and WT mice. Mice were dosed with saline during the dark period (ZT13), glucose (2g/kg; ip) in the dark period (ZT13), or glucose in the light period (ZT1; Fig 8A). Changes in ISF glucose and glutamate were assessed over the 11 hours following a peripheral glucose challenge. Most notably, time of day impacted the brain’s response to a metabolic challenge. During the dark period, ISF glucose increased in response to a systemic glucose challenge, but this effect was potentiated in the P301S mice compared to WT (Fig 8B, p<0.0001). Consistent with our SIRM data (Fig 6), ISF glutamate increased across the dark period post-glucose injection in the P301S mice compared to WT (Fig 8C, p<0.0001). These levels remained elevated for the duration of the dark period in P301S mice (Fig 8C) and coincided with a sharp decrease in ISF lactate availability, presumably due to increased lactate consumption (Suppl. Fig 5A). Together, these findings suggest that tau pathology enhances glucose metabolism to support increased glutamatergic tone during periods of activity. In contrast, during the light phase, systemic glucose raised ISF glucose to similar levels in both genotypes (Fig. 8D). However, ISF glutamate decreased in WT mice—likely reflecting reduced neuronal activity associated with quiet wakefulness or sleep—but remained elevated in P301S mice (Fig. 8G, p<0.0001), indicating sustained excitatory activity during the inactive phase. Together, these results suggest that tau pathology increases glutamatergic activity, which feeds forward to drive metabolic need and glucose utilization.

**Figure 8.**
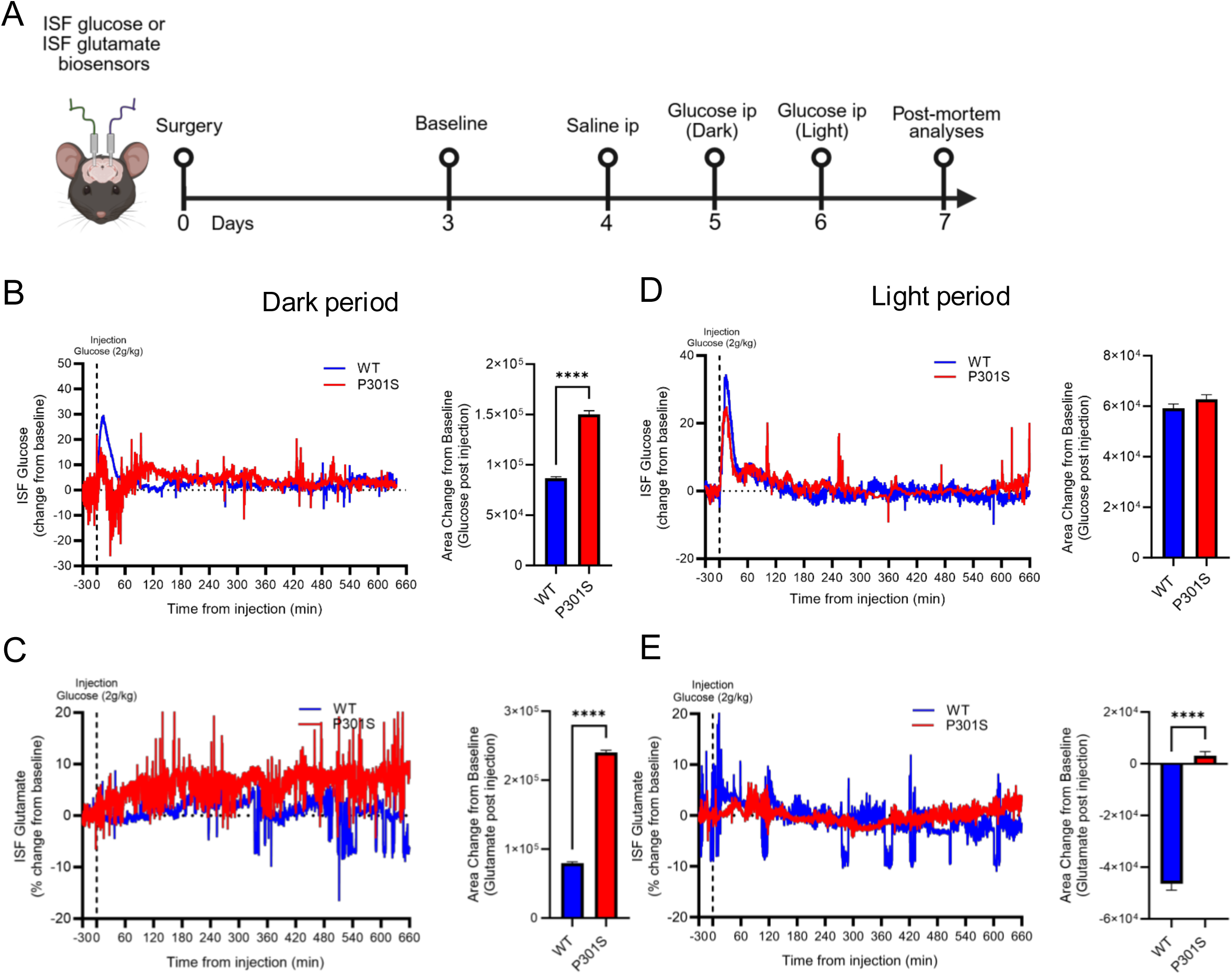
Glucose fuels ISF glutamate levels in P301S mice. (A) Experimental timeline for ISF glucose and glutamate biosensor recordings and i.p. glucose injections in 9 month P301S v WT mice. (B) During the dark period, ISF glucose is increased after glucose injection (ip) in P301S and WT brains. ISF glucose levels remain elevated in the in the P301S brain post-injection compared to WT. (C) ISF glutamate is increased in the P301S brain compared to WT following a glucose injection. (D) Levels of ISF glucose in P301S and WT brains are comparable after glucose injection during the light period. (E) ISF glutamate, however, decreased over the light period (or inactive period) in WT, where ISF glutamate remains above baseline in P301S mice, suggesting there is excess ISF glutamate in the P301S brain. Traces show 30 minutes before injection (baseline) and 11 hours post injection. ISF glucose and glutamate traces shown as percent change from baseline before and after glucose injection. Significance determined using area under the curve (net area change from baseline) and unpaired t-test. Data shown as means ± SEM. *n =* 3-4 mice/group. ***p<0.001, ****p<0.0001

## Discussion

Neuronal activity and cellular metabolism are fundamentally interdependent processes. Excitatory activity imposes high energetic demands on neurons requiring ATP production to fuel transmission, vesicle recycling, and neurotransmitter biosynthesis [52, 53]. Glucose is the primary substrate used to meet these energy needs and fluctuations in neuronal activity are tightly matched to glucose uptake and utilization [54, 55]. This metabolic-excitatory coupling is essential for maintaining network stability, plasticity, and cognitive function [56–58]. Disruption of this tightly regulated relationship, as seen in aging and disease, leads to inefficient energy utilization, synaptic dysfunction, and heightened vulnerability to metabolic stress [59, 60]. Tau pathology may sever this link by increasing neuronal excitability as hyperphosphorylated tau released from cerebrospinal fluid is shown to depolarize hippocampal neurons and elevate firing rates – resulting in elevated excitatory drive [35, 61]. Further, knockdown of tau reduces excitatory input and promotes inhibition, suggesting tau may play a role in tipping E/I balance towards excitotoxicity [62, 63]. As such, tau-induced hyperexcitability increases energy demand and induces metabolic inflexibility by forcing sustained glucose consumption to support excess neuronal activity and increase neurotransmitter synthesis.

This study provides new insight into the complex interplay between tau pathology and metabolism-excitability coupling. Our data demonstrates that tau pathology does not simply disrupt homeostasis, rather, it protects and preserves brain and whole-body metabolism against the effects of normal aging. Tau pathology, in particular hyperphosphorylated tau, preserves glucose tolerance, maintains circadian rhythms in interstitial fluid (ISF) glucose and lactate, and drives fuel preference for carbohydrates. Notably, these changes are associated with preserved synaptic mitochondrial function, despite region-specific cortical atrophy and cell loss. Moreover, tau pathology reshapes the brain’s metabolic and excitatory landscape – a relationship that is normally tightly coupled to maintain healthy brain function. Notably, we observed preservation of both whole-body and brain metabolism in P301S mice, which appeared to be driven by a highly excitable system. This was accompanied by increased glutamate enrichment and a concomitant reduction in GABAergic tone – indicating an excitatory/inhibitory (E/I) imbalance. These changes were dynamically modulated by time of day, suggesting a circadian component of tau-dependent alterations in metabolism-excitability coupling. Moreover, alterations in this coupling were not restricted to the brain but mirrored in whole body metabolism. These findings suggest that tau pathology induces a shift toward an excitatory, metabolically active state that may initially serve as a compensatory or protective role, but could later exacerbate vulnerability to neurodegeneration by creating a metabolically inflexible system.

The implications of these findings extend beyond tauopathies, such as Alzheimer’s disease (AD) and frontotemporal dementia. Hyperexcitability and metabolic remodeling are shared features across multiple neurological and systemic diseases, including epilepsy, type 2 diabetes (T2D), and sleep disorders [64–71]. For instance, disruptions in sleep architecture and circadian regulation are increasingly implicated in AD pathogenesis [72, 73]. Furthermore, hyperexcitability – a hallmark of early AD– may underlie the increased metabolic demand, elevated ISF glutamate, and altered lactate-glutamate coupling observed herein [31, 74–79]. The association between neurodegenerative diseases and metabolic disorders like T2D suggests that brain-derived changes in fuel utilization and neurotransmitter cycling may contribute to, or reflect, systemic metabolic shifts [60, 80]. Our findings highlight a potential mechanistic link through tau-dependent alterations in metabolic-excitatory coupling that could alter neuronal activity and therefore negatively impact sleep and metabolism associated with AD [81–84].

Perhaps these findings are not surprising when considering the physiological functions of normal tau. Tau is not merely a microtubule-associated protein; it plays important roles in regulating axonal transport, synaptic function, and neuronal excitability [38]. Recent evidence indicates under physiological conditions, tau is activity-dependent, released during high neuronal activity, and may facilitate adaptive responses to metabolic stress [37, 85–87]. In this context, the observed increase in glutamate biosynthesis could reflect a heightened metabolic demand to sustain excitatory transmission. The decreased expression of SynGO transcripts despite elevated glutamate flux may reflect structural degeneration paired with an overactive residual network attempting to compensate. This mismatch between metabolic supply and synaptic capacity could cause metabolic inflexibility caused by a maladaptive state of excitatory stress. Moreover, the shift towards a hyperexcitable and metabolically active state in tauopathy may reflect an aberrant amplification of tau’s role in supporting neuronal activation. However, while early-stage tau pathology may promote plasticity or compensation, the long-term consequences – particularly as tau becomes hyperphosphorylated and accumulates as neurofibrillary tangles – are likely deleterious [88–91]. Moreover, our findings suggest that tau-dependent hypermetabolism is linked to hyperphosphorylated tau itself rather than the effects of tau aggregation or neurodegeneration. Thus, these results could present an early target for AD prior to neuronal loss and cognitive impairment.

A central theme in our data is the disruption of E/I balance shifting toward excitation, as evidenced by increased glutamate and decreased GABA. This imbalance likely underpins the preserved ISF glutamate and decreased GABA levels. Reduced GABAergic tone alongside elevated glutamate suggests that tau pathology selectively impacts inhibitory circuits or neurotransmitter biosynthesis, consistent with prior reports [92–98]. Our findings suggest that this imbalance may also serve as a driver of metabolic preservation, where the energetic cost of sustaining excitation is met by glucose shunting to the brain and altered mitochondrial function. Importantly, the observed decrease in non-synaptic mitochondrial State V (succinate) suggests subtle impairments in oxidative capacity within non-neuronal or somatic compartments, while synaptic mitochondria remain intact. Notably, our findings do not support a model of primary metabolic failure. Rather, this supports a model of metabolic inflexibility – where resources are available but utilized inefficiently or constrained to specific cellular domains [46, 99–101]. These findings are consistent with reports showing that mitochondrial dysfunction in AD is compartmentalized and evolves gradually rather than catastrophically [102–107].

Our results also suggest that tau pathology imposes a unique form of metabolic resilience. Unlike age-matched controls, P301S mice retain glucose sensitivity, locomotor activity, and RER rhythmicity – all metrics that typically decline with age. This peripheral metabolic resilience is reflective of brain metabolic preservation, a phenomenon shown by us and others [108–111]. Moreover, this resilience may reflect tau-induced network excitability, which drives glucose utilization for biosynthetic pathways such as glutamate synthesis. The tau-dependent preservation of ISF rhythms supports the enhancement of circadian metabolic regulation. However, whether this remodeling is ultimately beneficial or contributes to accelerated neurodegeneration remains unclear. Moreover, elevated glutamate is neurotoxic in excess, and chronic excitability increases the risk of excitotoxic damage, oxidative stress, and network instability [21, 112–115].

Taken together, our results support a model in which tau pathology acts upstream of both excitability and energy metabolism. This may drive a feedforward cycle that may initially preserve function, but ultimately contributes to neurodegeneration. Whether the observed metabolic preservation is beneficial or harmful remains unclear – particularly in light of potential cell loss, reactive gliosis, or immune cell infiltration at later stages of disease. A key limitation of previous studies was the inability to distinguish between tau-driven alterations and secondary effects of neurodegeneration; however, our findings suggest that tau pathology itself promotes a hypermetabolic phenotype that preserves peripheral glucose clearance. Moreover, causality between E/I imbalance and metabolic remodeling remains unresolved: is hyperexcitability the cause or consequence of altered metabolism? Longitudinal studies, cell-type-specific interventions, and real-time metabolic imaging will be critical in disentangling these relationships.

In conclusion, our findings suggest that tau pathology reshapes excitability-metabolism coupling in a time-of-day-dependent manner. These changes reflect not only the pathological effects of tau but also its physiological role in regulating neuronal activity and metabolism. This dynamic interplay between excitation, metabolism, and circadian regulation offers a novel framework for understanding tauopathies and their comorbidities, including metabolic syndromes and sleep disruption. Targeting this feedforward excitatory-metabolic loop may hold promise for preserving function and delaying progression in tauopathies, such as Alzheimer’s disease.

## Data availability

The datasets used or analyzed in the current study are included within the manuscript.

## Acknowledgments

We would like to acknowledge Dr. Caitlin Carroll for her technical development on this project. We would like to acknowledge the Metabolic Phenotyping Shared Resource at Wake Forest University School of Medicine for the indirect calorimetry experiments. We would like to acknowledge the following grants: R01AG068330 (SLM), R01AG093847 (Macauley), BrightFocus Foundation A20201775S (SLM), Coins for Alzheimer’s Research Trust Grant (Macauley), P30AG072946 (SLM), R01AG060056 (LAJ), R01AG062550 (LAJ), R01AG080589 (LAJ), and the Alzheimer’s Association (LAJ). This research was supported by an Institutional Development Award (IDeA) from the NIGMS and NIH (P30GM127211) and the NIH Center of Biomedical Research Excellence (COBRE) in CNS Metabolism (CNS-Met; P20GM148326).

## Authors Contribution

SLM and REI conceived of the study. SLM, REI, DCL, PGS, and LAJ contributed to study design. REI, SMT, VGV, JBH, JL, HCW, and JAS performed experiments. REI, HCW, VGV, DCL, LAJ, and SLM performed data analysis and data interpretation. REI and SLM wrote the manuscript. All authors discussed the results and commented on the manuscript.

## Abbreviations

AD: Alzheimer’s disease
Aβ: amyloid beta
GCMS: Gas chromatography mass spectrometry
GTT: Glucose tolerance test
FAO: Fatty acid oxidation
FDG-PET: 2-Deoxy-2-[^18^F]fluoro-D-glucose positron emission tomography
ISF: Interstitial gluid
i.p.: Intraperitoneal
JTK: Jonckheere-Terpstra-Kendall
MCI: Mild cognitive impairment
OCR: Oxygen consumption rate
PBS: Phosphate buffered saline
PFA: Paraformaldehyde
Ptau: Hyperphosphorylated tau
P301S: P301S PS19
RER: Respiratory exchange ratio
TEE: Total energy expenditure
SIRM: Stable isotope resolved metabolomics
VCO2: Carbon dioxide production
VO2: Oxygen consumption
WT: Wildtype
4Rtg: Tau4RTg2652

## Supplementary Figures

**Suppl. Fig 1.**
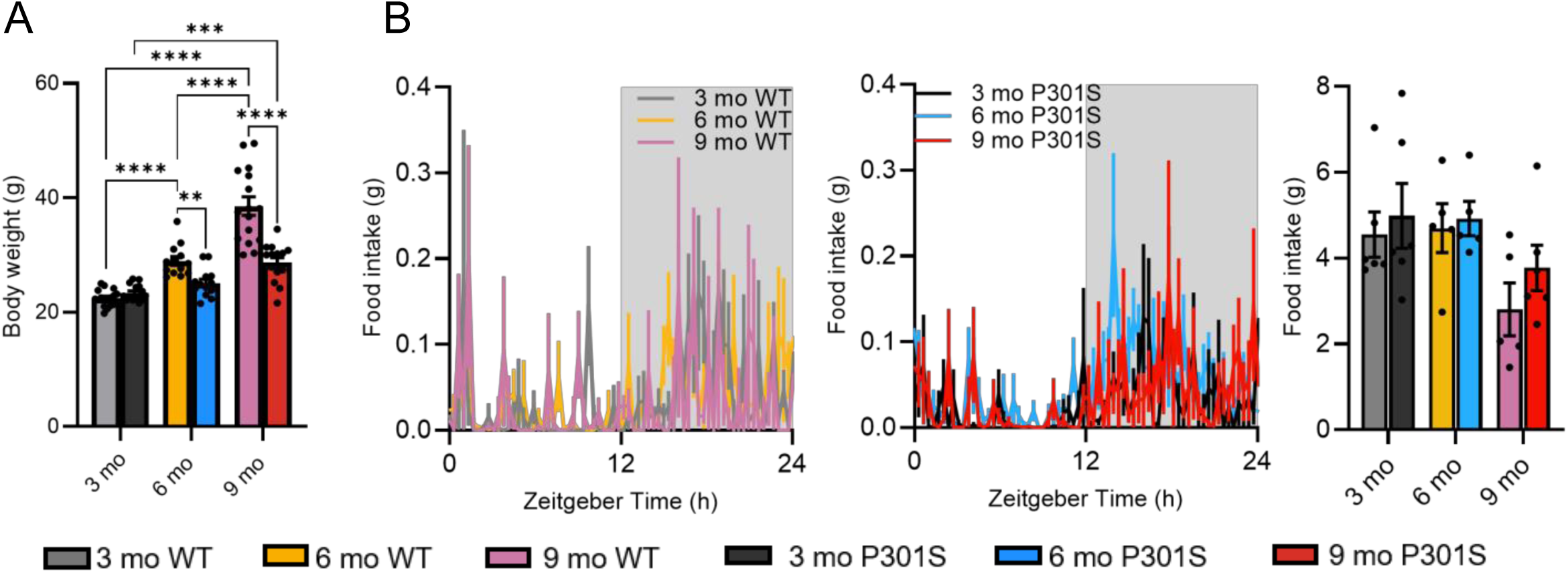
Tau pathology decreases body weight despite no change in food intake. (A) WT mice gain weight in a step wise fashion as they age from 3- to 6- to 9 months old. There is no difference in body weight between 3- and 6 month in P301S mice. By 9 months, P301S mice weigh less than age matched WT mice. (B) P301S and WT mice exhibit no change in food intake with age, despite differences in body weight. Statistical significance was determined using a two-way ANOVA with Tukey’s post-hoc tests. Data is represented by means ± SEM. *p<0.05, **p<0.01, ***p<0.001, ****p<0.0001

**Suppl. Fig 2.**
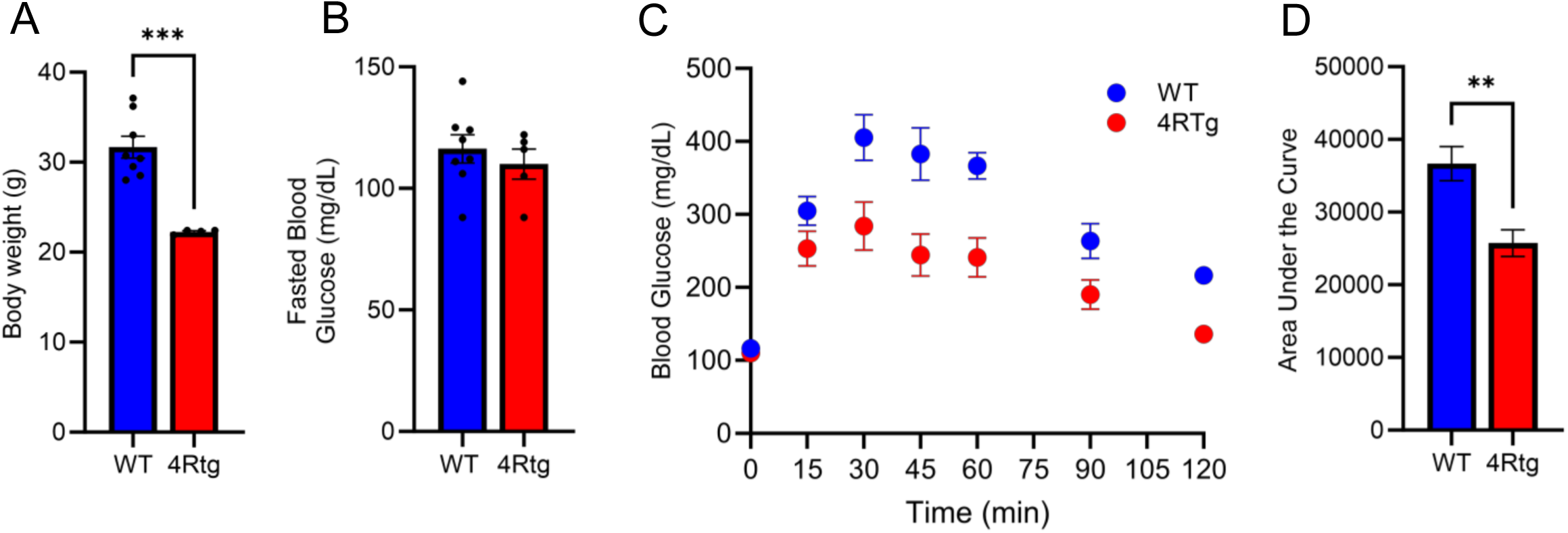
(A) 13 month 4RTg2652 mice weigh significantly less than age matched controls. (B) Fasting blood glucose levels were unchanged between 4Rtg and control mice. (C) 4RTg mice exhibited increased glucose tolerance compared to age matched controls as shown within the GTT curve by decreased peak and shorter tail. (D) Area Under the Curve indicated increased blood glucose levels in control mice compared to 4RTg mice. Significance determined by unpaired t -tests and two-way ANOVA with Sidak’s post-hoc corrections. *n* = 5-8 mice/group. Data is represented by means ± SEM. *p<0.05, **p<0.01, ***p<0.001

**Suppl. Fig 3.**
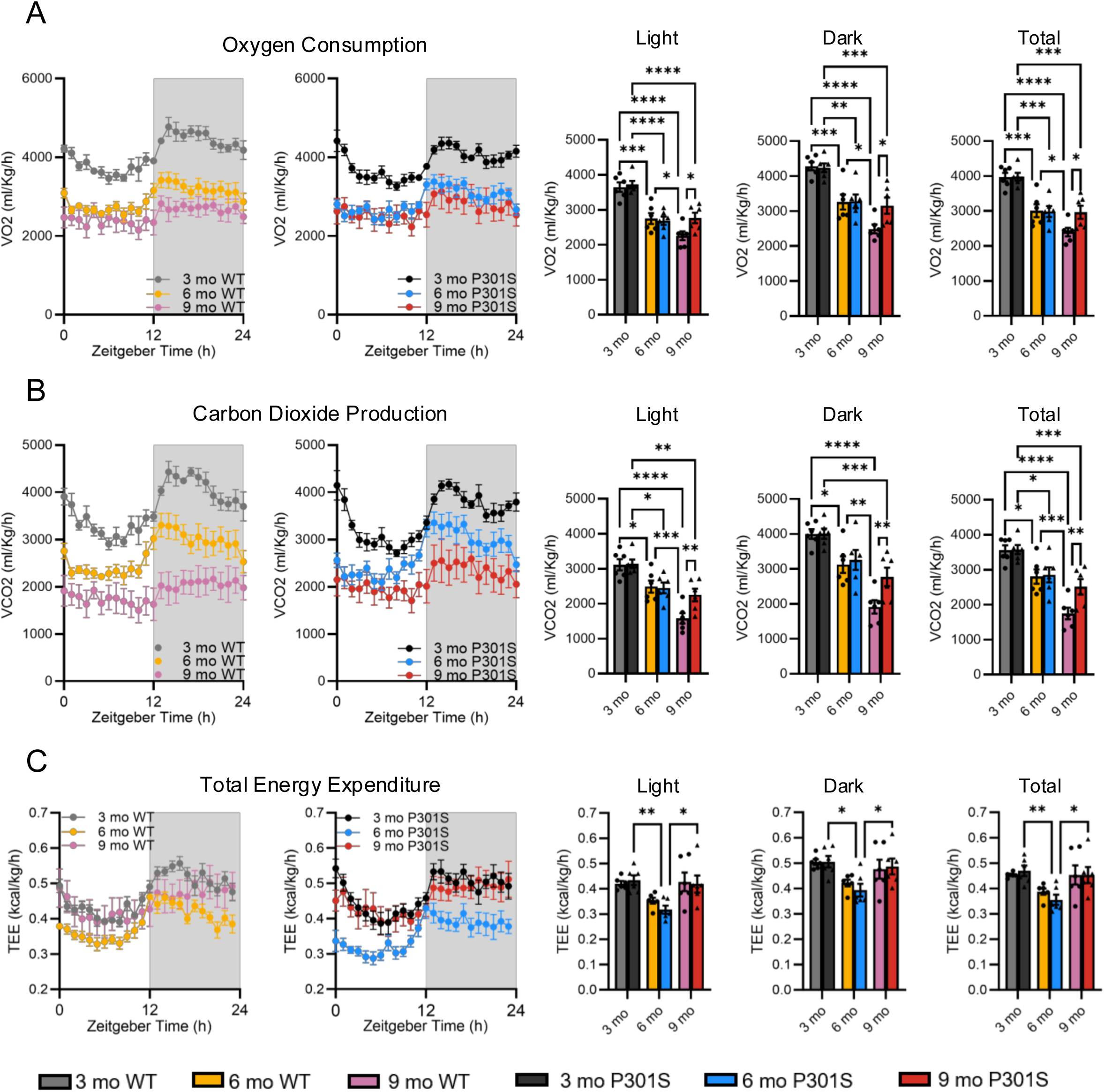
Tau pathology slows age dependent decreases in whole body metabolic characteristics. (A) As WT and P301S mice age, they exhibit decreased oxygen consumption (VO2) over the entire 24-hour period. However, 9-month-old P301S mice exhibit increased VO2 compared to WT mice. (B) As WT and P301S mice age, they exhibit decreased carbon dioxide production (CO2) over the entire 24-hour period. However, 9-month-old P301S mice exhibit increased VCO2 compared to WT mice. (C) At 6-months-old, P301S mice exhibit decreased total energy expenditure (TEE) compared to 3- and 9-month-old P301S mice at all times of day. Data reported as means ± SEM. *n =* 6 mice/group. Significance determined using two-way ANOVA with Tukey’s post-hoc correction. *p<0.05, **p<0.01, ***p<0.001, ****p<0.0001

**Supp. Table 1.** JTK Analysis for ISF glucose and lactate rhythmicity. (A) JTK Analysis confirmed the presence of diurnal ISF glucose rhythms in 3-, 6-, and 9-month-old P301S and WT mice. While ISF glucose rhythmicity is maintained in 9 mo WT mice, lag analysis suggesting the rhythm peaks during the light period. (B) JTK analysis confirmed diurnal ISF lactate rhythms are lost in 9 month WT mice but not P301S mice at any age. ADJ.P = adjusted p-value, Period = length of cycle, Lag = ZT time of peak, Amplitude = peak expression.

| Genotype | Age (mo) | ADJ.P | Period | Lag | Amp |
| --- | --- | --- | --- | --- | --- |
| WT | 3 | 3.17E-04 | 24 | 17 | 2.096 |
| WT | 6 | 1.09E-06 | 24 | 16 | 1.416 |
| WT | 9 | 5.5E-03 | 21 | 6 | 0.160 |
| P301S | 3 | 9.11E-06 | 24 | 15.5 | 1.237 |
| P301S | 6 | 5.79E-08 | 23 | 18 | 1.622 |
| P301S | 9 | 4.27E-03 | 24 | 19 | 0.918 |

**Supp. Table 1. JTK Analysis for ISF glucose and lactate rhythmicity.**
| Genotype | Age (mo) | ADJ.P | Period | Lag | Amp |
| --- | --- | --- | --- | --- | --- |
| WT | 3 | 7.98E-04 | 24 | 16.5 | 2.186 |
| WT | 6 | 1.09E-06 | 24 | 15.5 | 1.435 |
| WT | 9 | 5.7E-02 | 24 | 17 | 0.321 |
| P301S | 3 | 8.98E-10 | 24 | 16 | 1.481 |
| P301S | 6 | 4.03E-06 | 23 | 17.5 | 0.920 |
| P301S | 9 | 9.79E-08 | 24 | 17 | 0.918 |

**Suppl. Fig 4.**
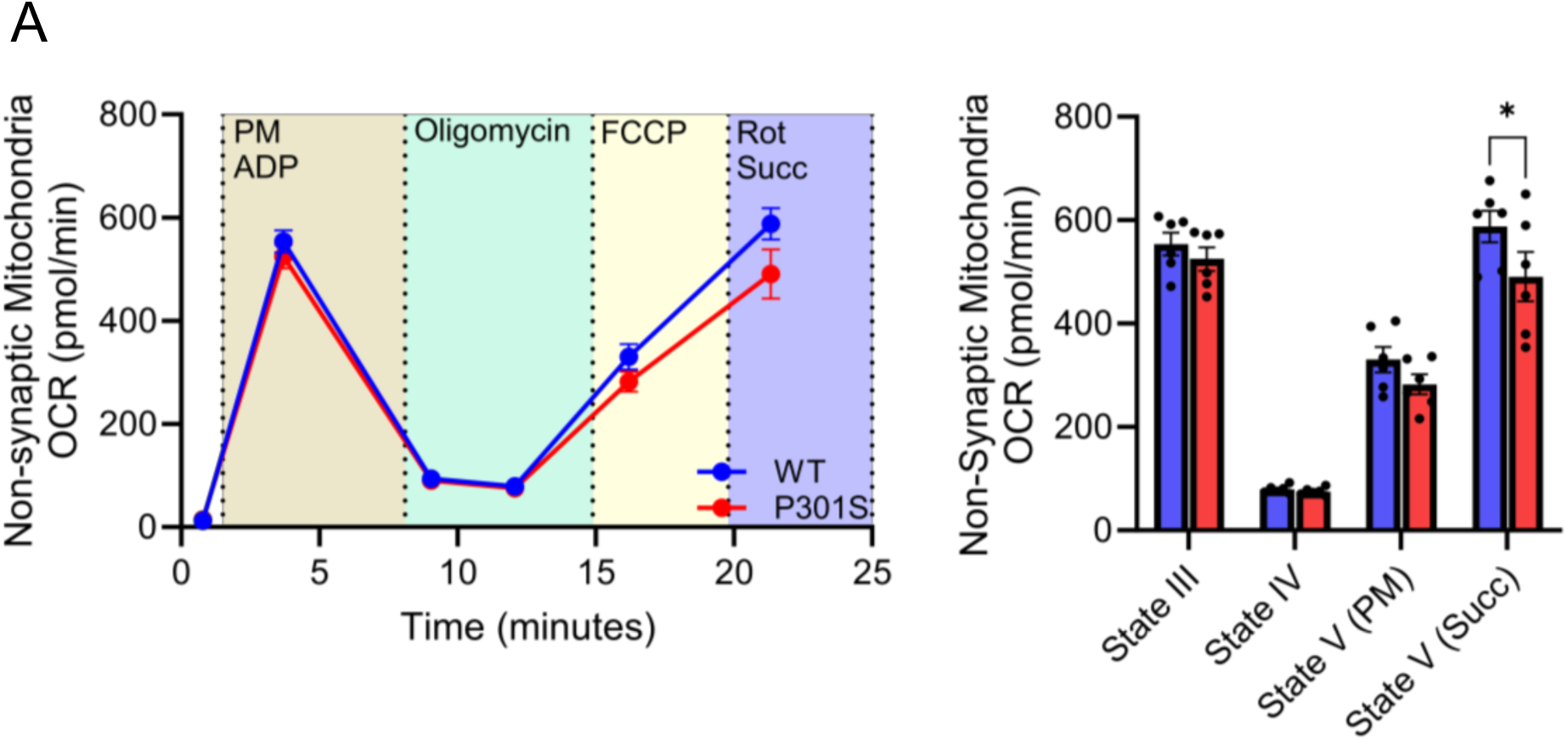
Tau pathology decreases non-synaptic mitochondrial State V respiration. (A) Oxygen consumption rate (OCR) is decreased in 9-month P301S mice in non-synaptic mitochondria, including mitochondria found in the neuronal soma and glia, during State V respiration. State V typically is associated with oxygen-limited respiration. Statistical significance was determined using a two-way ANOVA with Tukey’s post-hoc tests. *n =* 6 mice/group. Data is represented by means ± SEM. *p<0.05.

**Suppl. Fig 5.**
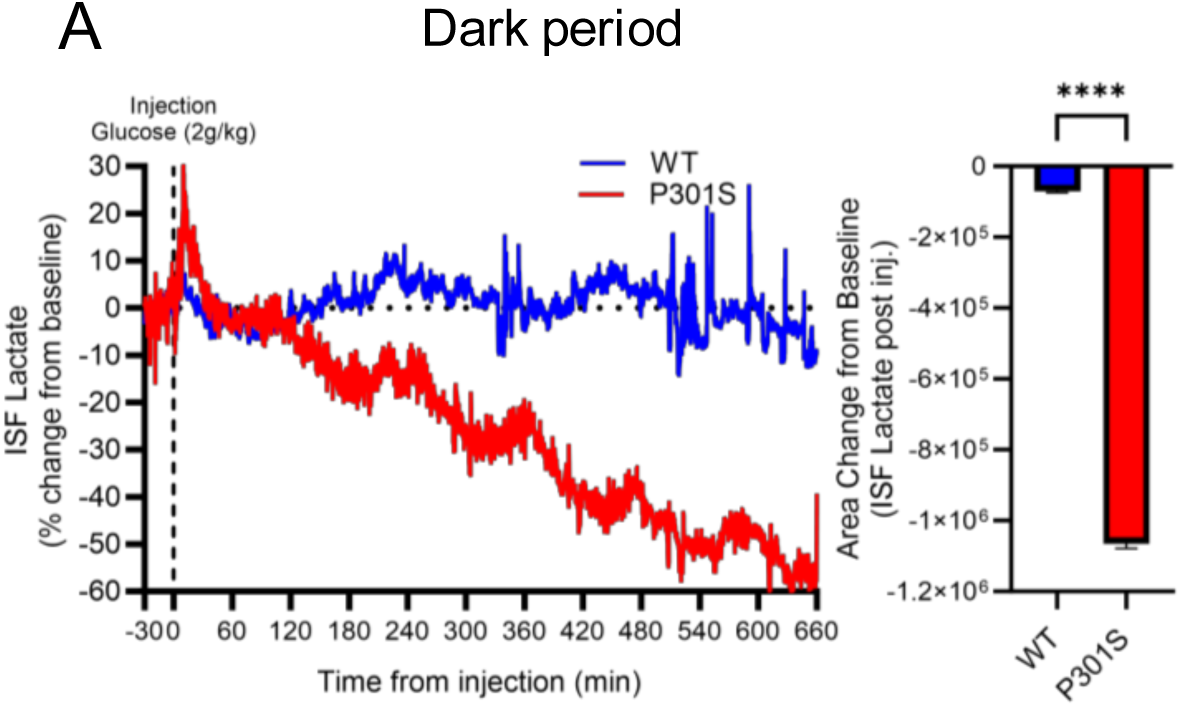
ISF lactate decreases following ip glucose challenge in the dark period in P301S mice, suggesting increased lactate consumption. (A) ISF lactate decreases in response to an ip glucose challenge in P301S mice, not WT mice, in the dark or active period. Significance determined using area under the curve (net area change from baseline) and unpaired t -test. Data shown as means ± SEM. *n =* 3-4 mice/group. ***p<0.001, ****p<0.0001

